# Bidirectional feedforward regulatory loop of Dicer-like 4 and flavonoids causes floral bicolor patterning in petunia and dahlia

**DOI:** 10.1101/2023.10.16.562536

**Authors:** Kazunori Kuriyama, Sho Ohno, Niichi Yamazaki, Midori Tabara, Hisashi Koiwa, Hiromitsu Moriyam, Toshiyuki Fukuhara

**Affiliations:** Department of Applied Biological Sciences, Tokyo University of Agriculture and Technology, 3-5-8 Saiwaicho, Fuchu, Tokyo 183-8509, Japan; Graduate School of Agriculture, Kyoto University, Kyoto, Kyoto 606-8502, Japan; Ritsumeikan-Global Inovation Research Organization, Ritsumeikan University, 1-1-1, Noji-Higashi, Kusatsu, Shiga, 525-8577, Japan; Vegetable and Fruit Improvement Center and Department of Horticultural Sciences, Texas A&M University, College Station, TX 77843, USA; Institute of Global Innovation Research, Tokyo University of Agriculture and Technology, 3-5-8 Saiwaicho, Fuchu, Tokyo 183-8509, Japan

**Author notes:** Corresponding author: Toshiyuki Fukuhara, Tokyo University of Agriculture and Technology, E-mail.

**Keywords:** Bicolor flower, DCL4, Feedforward regulation, Flavonoid, RNAi

## Abstract

Floral bicolor pigmentation is caused by naturally occurring RNA interference (RNAi) in some cultivars of petunia and dahlia. In both plants, the chalcone synthase gene is highly expressed only in the pigmented region of bicolor petals. However, it remains unknown why RNAi is induced only in the unpigmented region. To elucidate the mechanism of this clear bicolor pattern formation, we examined the dicing activity of Dicer-like 4 (DCL4), which produces small interfering RNAs essential for the induction of RNAi. We showed that the crude extract in the pigmented petal region inhibits dicing activity of DCL4, but not when flavonoids were depleted from the extract. Moreover, we showed the inhibitory activity was associated with flavonoid aglycons. The *in vivo* dicing activities were detected in the intact protoplasts prepared from the unpigmented region but not from the pigmented region. These results suggest that in the unpigmented region, flavonoids that inhibit DCL4 are not synthesized, and RNAi is maintained, whereas in the pigmented region, DCL4 is inhibited by flavonoids, RNAi is not induced, and anthocyanin biosynthesis is maintained, which ensures RNAi inhibition. Therefore, a clear bicolor pattern is generated by the bidirectional feedforward mechanism of antagonizing DCL4 and flavonoids.

## Introduction

In eukaryotes, RNA interference (RNAi), also known as RNA silencing, is a mechanism for regulating gene expression triggered by double-stranded RNAs (dsRNAs) and/or small interfering RNA (siRNA) derived from them (Baulcombe, 2004). RNAi is roughly divided into transcriptional gene silencing (TGS), which regulates gene expression during mRNA biogenesis, and post-transcriptional gene silencing (PTGS), which regulates gene expression by mRNA degradation (Borges and Martienssen, 2015). Dicer-like 4 (DCL4) cleaves long dsRNAs into 21-nt siRNAs that function in PTGS, while Dicer-like 3 (DCL3) cleaves short dsRNAs into 24-nt siRNAs that are involved in TGS (Nagano et al., 2014). RNAi (PTGS) was discovered approximately three decades ago when the exogenous chalcone synthase (CHS) gene, a key enzyme for anthocyanin biosynthesis, was overexpressed in transgenic petunia (*Petunia hybrida*) plants to produce dark colored flowers, although white flowers were unexpectedly observed (Napoli et al., 1990; van der Krol., 1990). In these transgenic petunia plants, RNAi was triggered by abnormal CHS transcripts during overexpression of the exogenous CHS gene, suppressing both exogenous and endogenous CHS genes (also known as co-suppression). Among non-transgenic garden petunia cultivars, two bicolor strains, the ‘Star’ strain, in which the unpigmented (white) regions are formed along the veins of the corolla as a star shape, and the ‘Picotee’ strain, in which the outer edge of the corolla is unpigmented (white), are well known (Figure 1). Both strains have the CHS genes in tandem on the same chromosome during breeding, and these genes may produce abnormal CHS transcripts and induce RNAi against the CHS genes (Koseki et al., 2005; Griesbach et al., 2007; Morita et al., 2012; Kasai et al., 2013).

**Figure 1.**
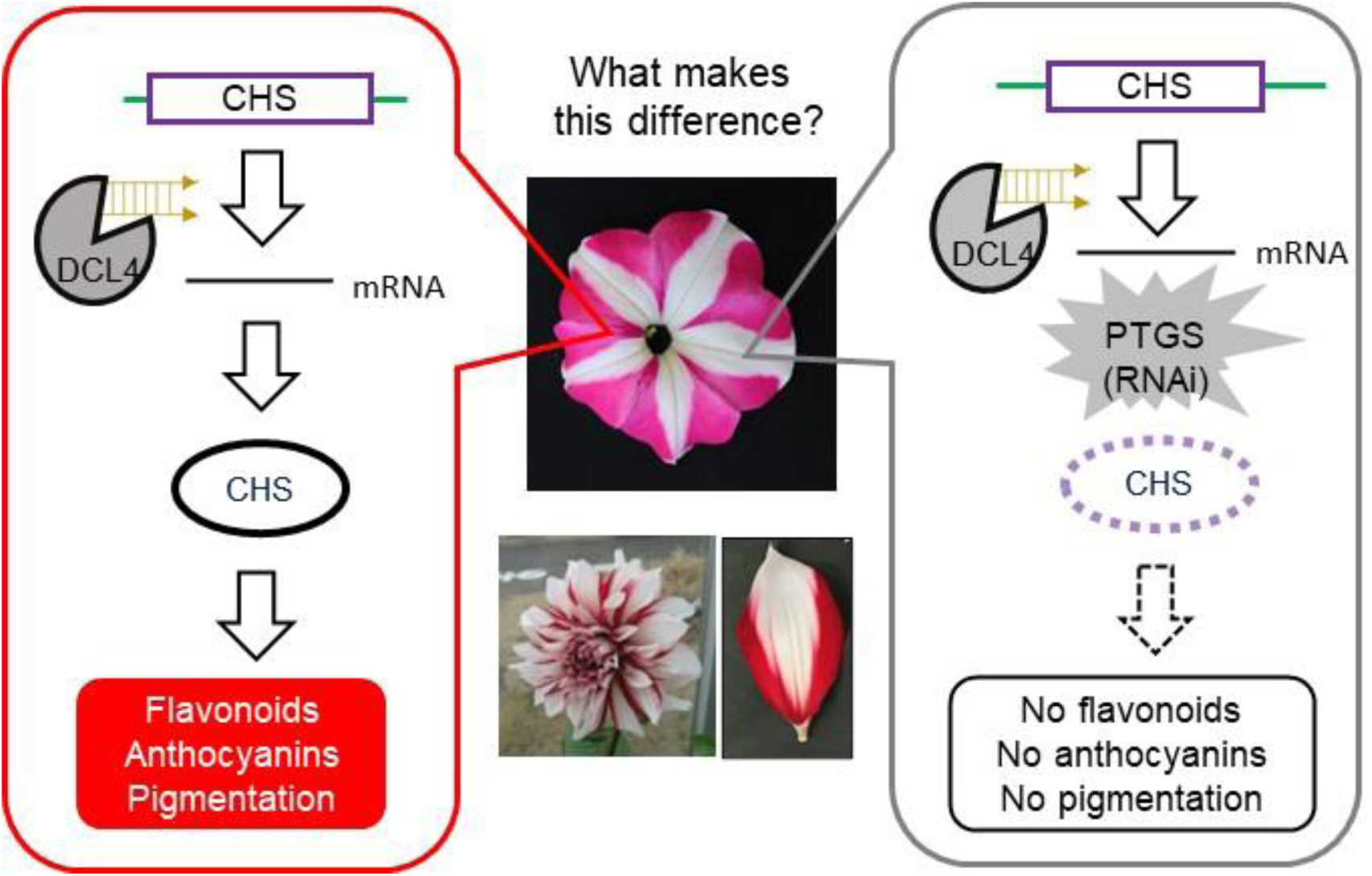
Bicolor flowers of petunia and dahlia and a schematic drawing of naturally occurring RNAi in bicolor petals. Bicolor flowers of cv. ‘Yuino’ (dahlia) and cv. ‘Rondo Rose Star’ (petunia) are shown. RNAi is induced only in the unpigmented regions of bicolor flowers of both plants. It remains unknown why the RNAi-induced unpigmented region and the RNAi-uninduced pigmented region coexist within the same petal.

Dahlia (*Dahlia variabilis*), a horticultural plant with over 50,000 cultivars created in the last 100 years, has flowers of various colors, sizes, and shapes (McClaren, 2009). Floral pigments, such as anthocyanins, flavonoids, and butein, also differ considerably among cultivars (Deguchi et al., 2013). One of the most attractive dahlia cultivars is the bicolor cultivar, in which the petals are clearly divided into pigmented bases (peripheral region) and unpigmented tips (central region) (Figure 1). It is known that RNAi for the CHS gene occurs only in the unpigmented region of bicolor petals in cultivar (cv.) ‘Yuino’ (Figure 1). The reason for this RNAi induction against the CHS gene is thought to be that modern dahlia cultivars have an octoploid genome during breeding, namely an increased gene dosage (Ohno et al., 2011). Bicolor cultivars induced by naturally occurring RNAi have been reported not only for petunia and dahlia, but also for gentian and bougainvillea (Ohta et al., 2021; Ohno et al., 2021).

Although it has been reported that these bicolor flowers of various ornamental plant species are caused by RNAi, it remains unknown why the RNAi-induced region and the RNAi-uninduced region coexist within the same petal (Figure 1). Since all cells in both regions have the same genome, the entire petal should not be pigmented by RNAi. Within the same tissue (petal), RNAi is induced in only one region where CHS gene expression is completely knocked down, but RNAi is not induced in the other region where the CHS gene strongly expressed (Figure 1). This spatial control mechanism of RNAi induction has never been elucidated. In many cases of naturally occurring RNAi reported so far, the target gene for RNAi encodes enzymes responsible for anthocyanin biosynthesis (e.g., CHS), and pigment biosynthesis is inhibited by RNAi, resulting in the appearance of unpigmented (white) regions. We conceived that inhibitory metabolites, such as anthocyanins or related secondary metabolites, may be involved in the spatial control of RNAi. Recently, we also reported that the enzyme activity of DCL4, an essential enzyme for RNAi, is regulated under various conditions (Seta et al., 2017; Tabara et al., 2023). Based on a model of inhibitory metabolites that target DCL4 activity to suppress RNAi in a spatial manner (Tabara et al., 2023), we hypothesized that DCL4 is activated in the unpigmented regions of bicolor flower petals, and pigment biosynthesis is subsequently inhibited by RNAi. On the other hand, in the pigmented region, the pigment or its related metabolites inhibit DCL4 activity, and RNAi is no longer induced, resulting to maintain anthocyanin biosynthesis.

In this study, we investigated bicolor cultivars of petunia and dahlia to verify this hypothesis. DCL4 dicing activity was detected in the unpigmented region of bicolor petals of dahlia and petunia but not in the pigmented region, as was hypothesized. In addition, pigmented regions of these bicolor petals contained large amounts of phenolic compounds such as flavonoids including anthocyanins, and the DCL4 activity was recovered by removing phenolic compounds by polyvinylpolypyrrolidone (PVPP) from enzymatic fractions prepared from the pigmented regions. In bicolor petals of dahlia and petunia, flavonoids that inhibit dicing activity are not biosynthesized in the unpigmented region where RNAi is stably maintained, whereas flavonoids inhibit DCL4 activity, namely RNAi, and maintain their biosynthesis in the pigmented region by themselves. Therefore, a clear bicolor pattern is generated by this bidirectional feedforward control mechanism composed of DCL4 and flavonoids.

## Results

### Dicing activity was detected only in the unpigmented region of bicolor petals

To examine if floral pigmentation patterns correlate with the presence or absence of DCL4 activity, we performed the dicing (Dicer) assay using cell-free extracts prepared from the dissected flower petals. The ^32^P-labelled 50 nt dsRNA was used as the substrate, and 21-, 24- and 31-nt RNA fragments were detected as products of putative DCL3 and DCL4 (a detailed explanation of the dicing assay is shown in Supplemental Figure S1; Fukudome et al., 2011; Nagano et al., 2014; Tabara et al., 2023). When cell-free extracts prepared from the pigmented (red) and unpigmented (white) regions of petals in the bicolor dahlia cv. Yuino were used as an enzyme fraction, dicing products of 21-nt, 24-nt and 31-nt RNA were detected only in reaction mixtures using the enzyme fractions from the unpigmented region (Figure 2A and Supplemental Figures S2A and S3), coinciding with the active RNAi against the CHS gene even though the expression levels of DCL4 were similar in both pigmented and unpigmented regions (Supplemental Figure S4). In the immature flower buds, in which bicolor pattern (pigmentation) had not developed yet, DCL4 activity was detected in both peripheral and central regions of young petals (Figure 2A). Namely, the presence or absence of DCL4 activity was consistent with the presence or absence of pigments (i.e., the presence or absence of RNAi against the CHS gene) rather than the regions in the petals. Therefore, the regulation of DCL4 activity is likely involved in the bicolor pattern formation in the bicolor flowers of dahlia. Given that a difference in DCL4 activity develops as pigmentation occurs, the subtle difference between the two regions, where anthocyanins accumulate and not accumulate, may trigger the production of a clear bicolor pattern (Figure 2A). These petals do not exhibit an intermediate color (pink).

**Figure 2.**
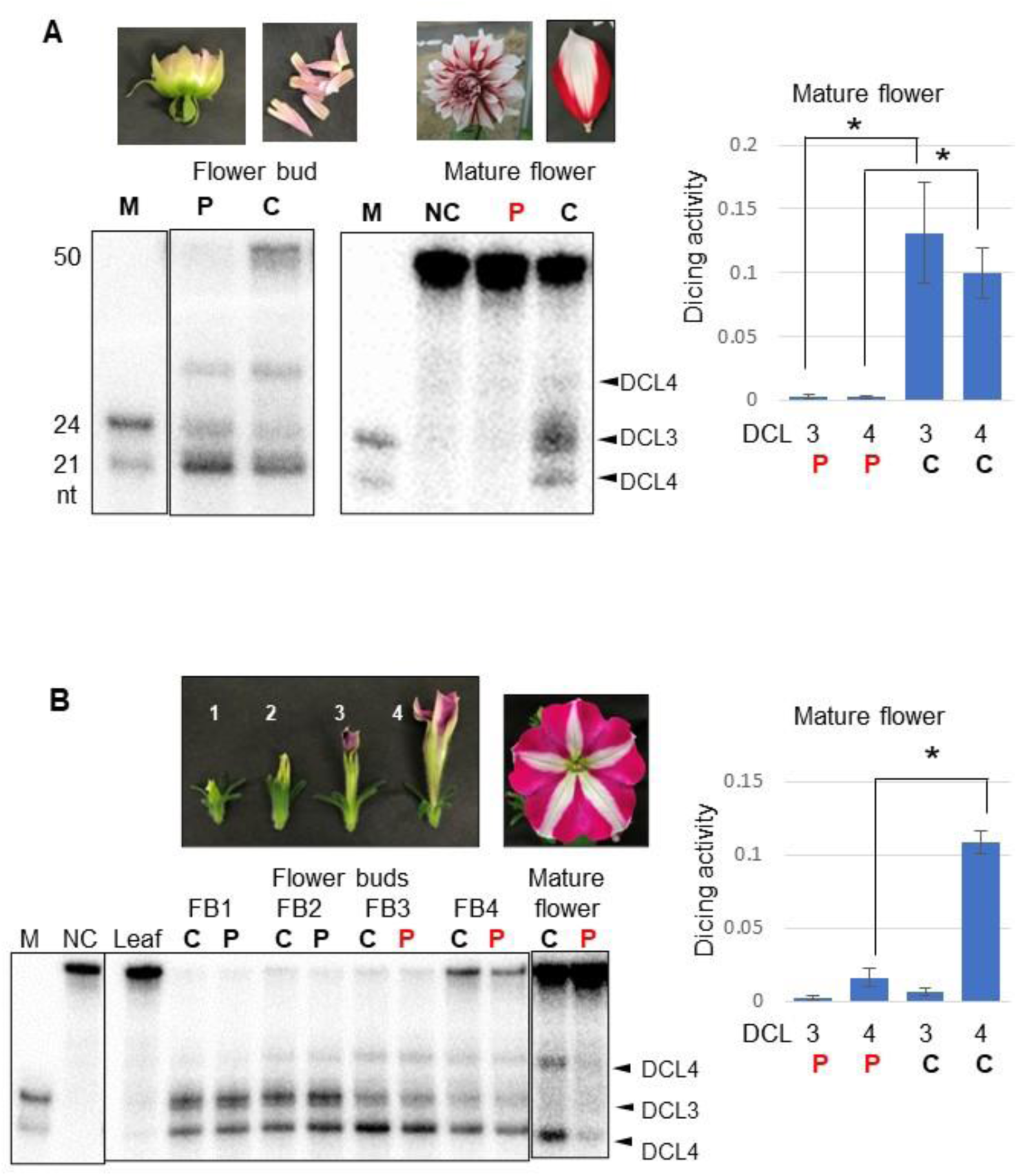
Dicing activity was detected only in the unpigmented region of dahlia and petunia bicolor petals. The dicing activities producing 21-nt, 24-nt and 31-nt RNAs from ^32^P-labeled 50-nt dsRNA with blunt ends as a substrate were detected from crude extracts prepared from petals of flower buds (FB) and mature flowers in dahlia cv. Yuino (**A**) and petunia cv. Rondo Rose Star (**B**). Cleaved RNA products of 21 and 31 nt were assumed to be produced by DCL4, and that of 24 nt was assumed to be produced by DCL3 (see Supplemental Figure S1). Autoradiographs of cleavage products analyzed by denaturing 15% PAGE are shown. Lanes C and P indicate the central and peripheral regions, respectively, in petals, and red lane P indicates the pigmented peripheral region in petals. Lanes M and NC indicate molecular weight markers of 21-nt and 24-nt ssRNAs and negative control (no crude extracts added), respectively. Error bars indicate the standard errors of six (**A**) and three (**B**) biological replicates, respectively (see Supplemental Figures S2 and S4). Black asterisks indicate significant differences between the central (C) and peripheral (P) regions using a Student’s test (p<0.01).

High dicing activities derived from putative DCL3 and DCL4 were also detected in both pigmented and unpigmented regions of bicolor petals of petunia flower buds (cv. Rondo Rose Star, Figure 2B and Supplemental Figures S2B). In petals of mature petunia flowers, however, the DCL4 activity producing 21-nt and 31-nt RNA was detected only in the unpigmented region but not in the pigmented region (Figure 2B and Supplemental Figures S2B and S5). Therefore, in both dahlia and petunia, DCL4 activity, which is essential for RNAi induction, was detected only in the unpigmented region where RNAi occurred. The spatial induction of RNAi was consistent with the spatial maintenance of DCL4 activity (Figure 1 and Supplemental Figures S2, S3, and S5).

### Detection of siRNAs derived from the CHS genes in the unpigmented region of bicolor petals

siRNAs derived from the CHS gene occur in the unpigmented regions of bicolor petals in petunia and dahlia (Koseki et al., 2005; Ohno et al., 2011; Morita et al., 2012). In dahlia, *DvCHS2*-derived siRNA accumulated in the unpigmented region of bicolor petals (Figure 3A), where DCL4 activity was detected (Figure 2A). Also in petunia, *PhCHS*-A-derived siRNAs were present from the early stage of bud development, and siRNAs were detected only in the unpigmented regions of bicolor petals (Figure 3B and Supplemental Figure S6), where DCL4 activity was detected (Figure 2B). These results support the hypothesis that the difference in DCL4 activity defines the region where RNAi is induced.

**Figure 3.**
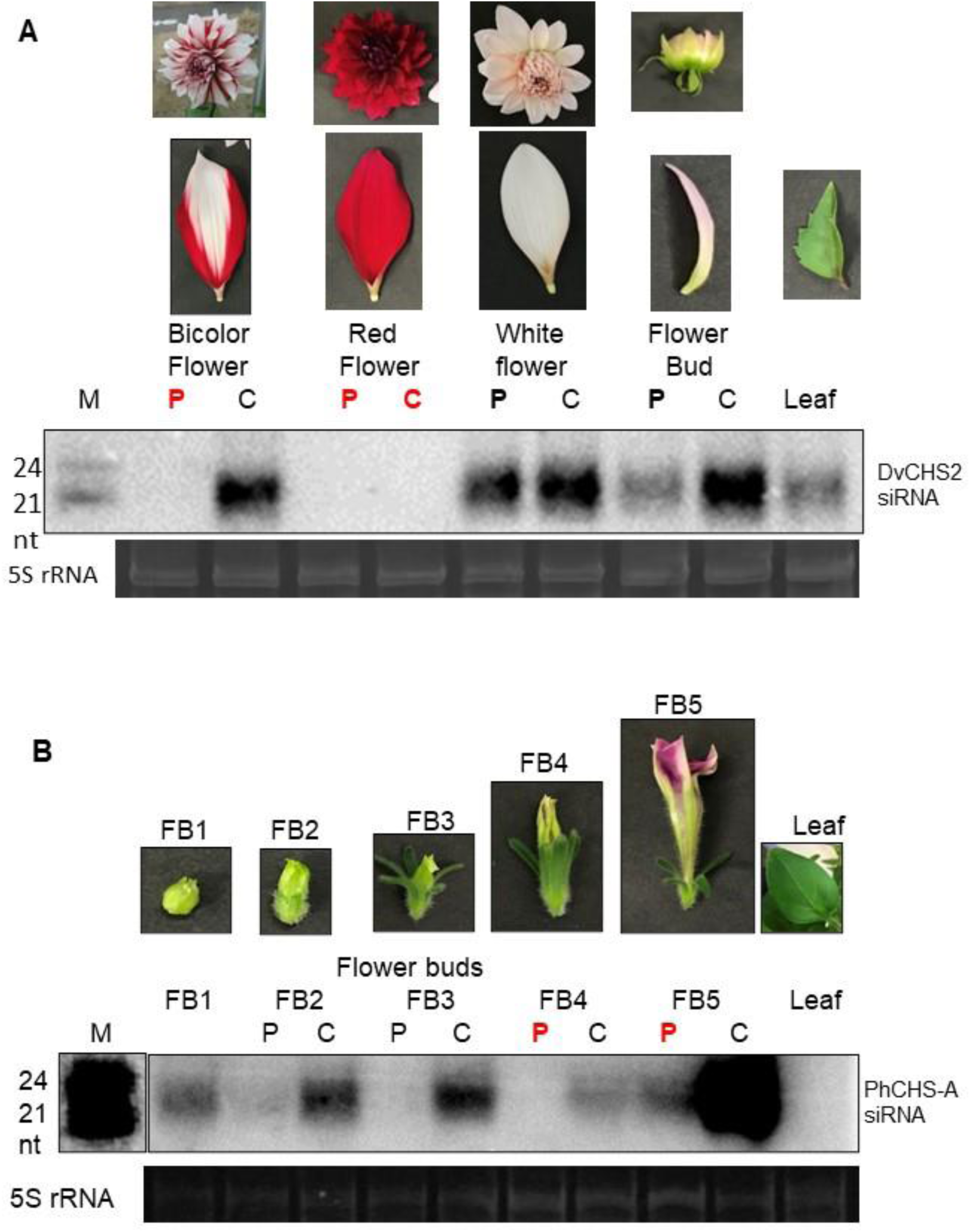
Detection of siRNAs derived from the CHS genes in the unpigmented regions of bicolor petals. siRNAs derived from the CHS gene in dahlia cv. Yuino (**A**) and petunia cv. Rondo Rose Star (**B**) were detected by Northern hybridization using ^32^P-labeled DNA fragments derived from the DvCHS2 (**A**) and PhCHS-A (**B**), respectively, as probes. Lanes C and P indicate the central and peripheral regions, respectively, in bicolor petals, and lanes red C and P indicates the pigmented regions. M indicates molecular weight markers of 21-nt and 24-nt ssRNAs, and rRNA indicates rRNA bands stained by ethidium bromide as a loading control.

### Putative inhibitors for DCL3 and DCL4 accumulated in the pigmented region of bicolor petals

In bicolor petals of dahlia, the dicing activities of DCL3 and DCL4, which produce 24-nt and 21-nt siRNAs, respectively (see Supplemental Figure S1), were detected in the unpigmented region but not in the pigmented region. We hypothesized that some metabolites inhibitory for DCL3/DCL4 accumulated in the pigmented region of bicolor petals, similar to the DCL3/DCL4 inhibitors in soybean leaves (Tabara et al., 2023). To verify this hypothesis, we examined the dicing activity of cell-free extracts prepared from the unpigmented region when the crude extracts prepared from the pigmented region were added. The dicing activity in the unpigmented region was strongly inhibited by the crude extracts prepared from the pigmented region of bicolor petals (Figure 4), indicating that the cell-free extracts prepared from the pigmented region of bicolor petals contained putative inhibitors of DCL3 and DCL4.

**Figure 4.**
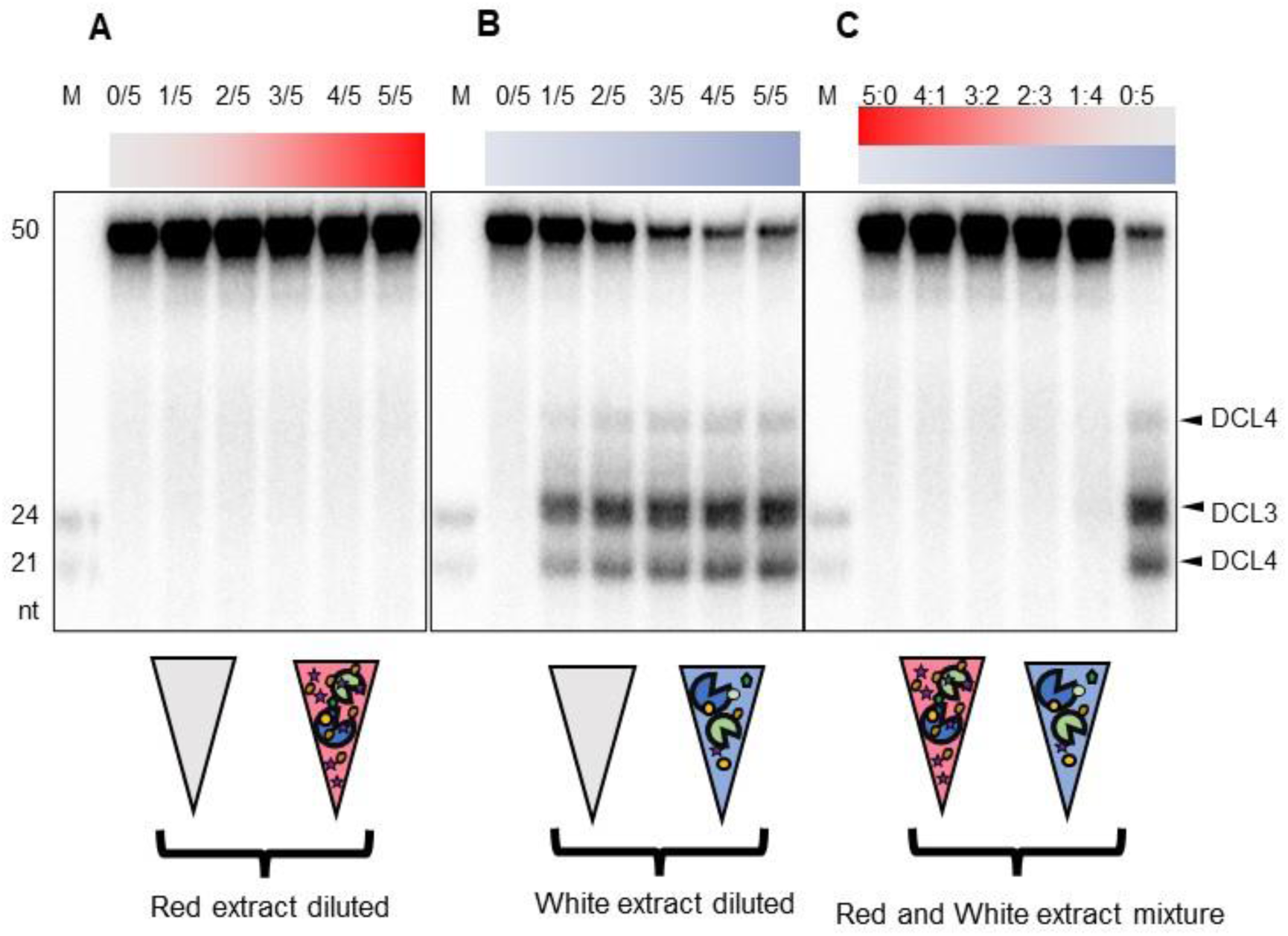
Putative inhibitors for the dicing activity of DCL3 and DCL4 accumulated in the pigmented region of bicolor dahlia petals. Dicing assays were performed using crude extracts prepared from the pigmented and unpigmented regions of bicolor dahlia petals as enzyme fractions. Pigmented crude extracts diluted with an extraction buffer (**A**), unpigmented crude extracts diluted with an extraction buffer (**B**), and mixtures of pigmented and unpigmented crude extracts (**C**) were used as enzyme fractions. Autoradiographs of cleaved RNA products of 21, 24, and 31 nt from ^32^P-labeled 50-nt dsRNA analyzed by denaturing 15% PAGE are shown (see Supplemental Figure S1). M indicates molecular weight markers of 21-nt and 24-nt ssRNAs.

### Flavonoids, which were absorbed by PVPP, accumulated more in the pigmented region of bicolor petals

Based on our previous findings (Tabara et al., 2023), we predicted that the inhibitory substances for the dicing activities are phenolic compounds, such as flavonoids. Therefore, we quantified total phenolic compounds in bicolor petals of dahlia and petunia by the Folin-Ciocalteu method (Makker et al., 1993; FAO/IAEA, 2000), and prepared flavonoid-depleted crude extract for analysis using PVPP, which specifically absorbs flavonoids (Figure 5A, Doner et al., 1993). The amount of total phenolic compounds was determined before and after PVPP treatment in petals of the unpigmented flower buds, pigmented flower buds, and mature flowers of dahlia and petunia. The pigmented regions of bicolor petals of both dahlia and petunia contained high levels of phenolics, and approximately 50-80% of total phenolics was absorbed by PVPP (Figure 5B). Using high-performance liquid chromatography, we identified apigenin and pelargonidin as the major phenolics in the pigmented region of bicolor dahlia petals (Figure 5C), consistent with a previous report (Ohno et al., 2018). Notably, most flavonoids that accumulated in the pigmented region were absorbed by PVPP (Figure 5C). In the case of petunia, the major flavonoid in the pigmented region of bicolor petals was peonidin (Figure 5D). Therefore, these results indicate that phenolic compounds including flavonoids, which may inhibit DCLs, highly accumulated in the pigmented region of bicolor petals of dahlia and petunia.

**Figure 5.**
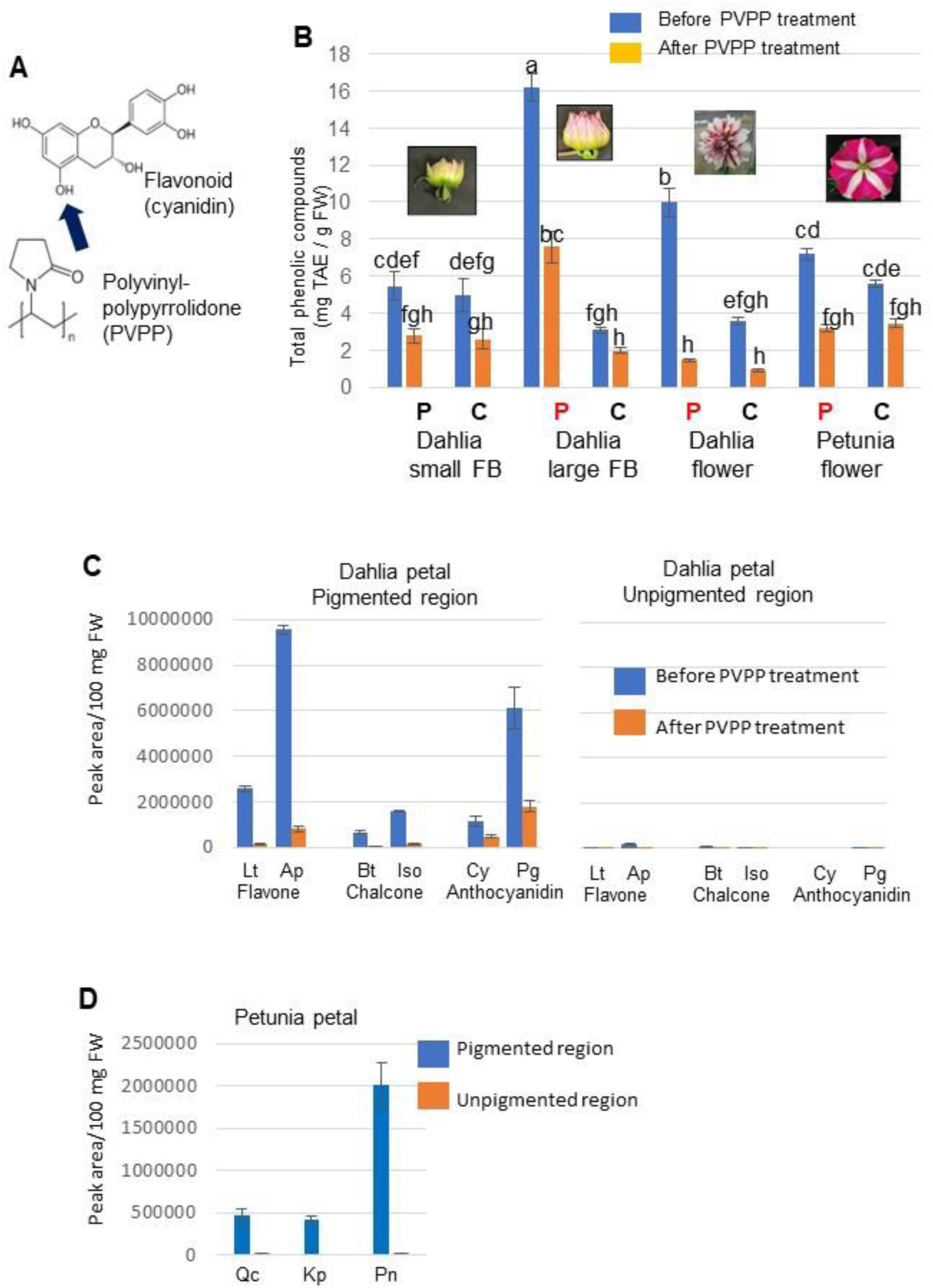
Identification and quantification of phenolic compounds that accumulated in bicolor petals of dahlia and petunia. **A**) Structural formulas of flavonoid (cyanidin) and polyvinylpolypyrrolidone (PVPP). **B**) Petals of unpigmented flower buds (small FB), pigmented flower buds (large FB), and mature bicolor flowers (flower) of dahlia (cv. Yuino) and petunia (cv. Rondo Rose Star) were divided into peripheral (P) and central (C) regions, from which total phenolic compounds were extracted. Half of the extracts were treated with PVPP. Total amount of phenolic compounds before and after PVPP treatment was quantified by the Folin-Ciocalteu method, and the quantitative value was based on tannic acid (Tannic Acid Equivalent: TAE). Three replicates were performed, and bars indicate standard errors. Different letters indicate significant differences using a Tukey’s test (p<0.05). **C**) Identification and quantification of flavonoids in the pigmented (left) and unpigmented (right) regions of bicolor dahlia petals. Extracts from dahlia petals before and after PVPP treatment were analyzed by HPLC. Lt, luteolin; Ap, apigenin; Bt, butein; Iso, isoliquiritigenin; Cy, cyanidin; Pg, pelargonidin. **D**) Identification and quantification of flavonoids in the pigmented and unpigmented regions of bicolor petunia petals. Extracts from petunia petals were analyzed by HPLC. Qc, quercetin; Kp, kaempferol; Pn, peonidin.

### PVPP removed Dicer inhibitors from crude extracts and restored dicing activity

By adding PVPP to the extraction buffer, we were able to prepare an enzyme fraction from which most flavonoids were removed (Figure 6). In both dahlia (Figure 6A) and petunia (Figure 6B and Supplemental Figure S7), high dicing activity equivalent to that detected in the unpigmented region was detected in the PVPP-treated fraction derived from the pigmented region. The addition of PVPP to the enzyme fraction derived from the unpigmented region did not affect the dicing activity (Figures 6A and 6B, lanes W). These results indicate that the inhibitors for DCL3 and DCL4 activities were phenolic compounds, probably flavonoids, that can bind to PVPP. Furthermore, the levels of both DCL3 and DCL4 proteins were similar in both pigmented and unpigmented tissues in the bicolor petals. These results indicate that flavonoids including anthocyanins highly accumulate in the pigmented region of bicolor petals and inhibit the dicing activities of DCL3 and DCL4 (Figures 2, 4-6).

**Figure 6.**
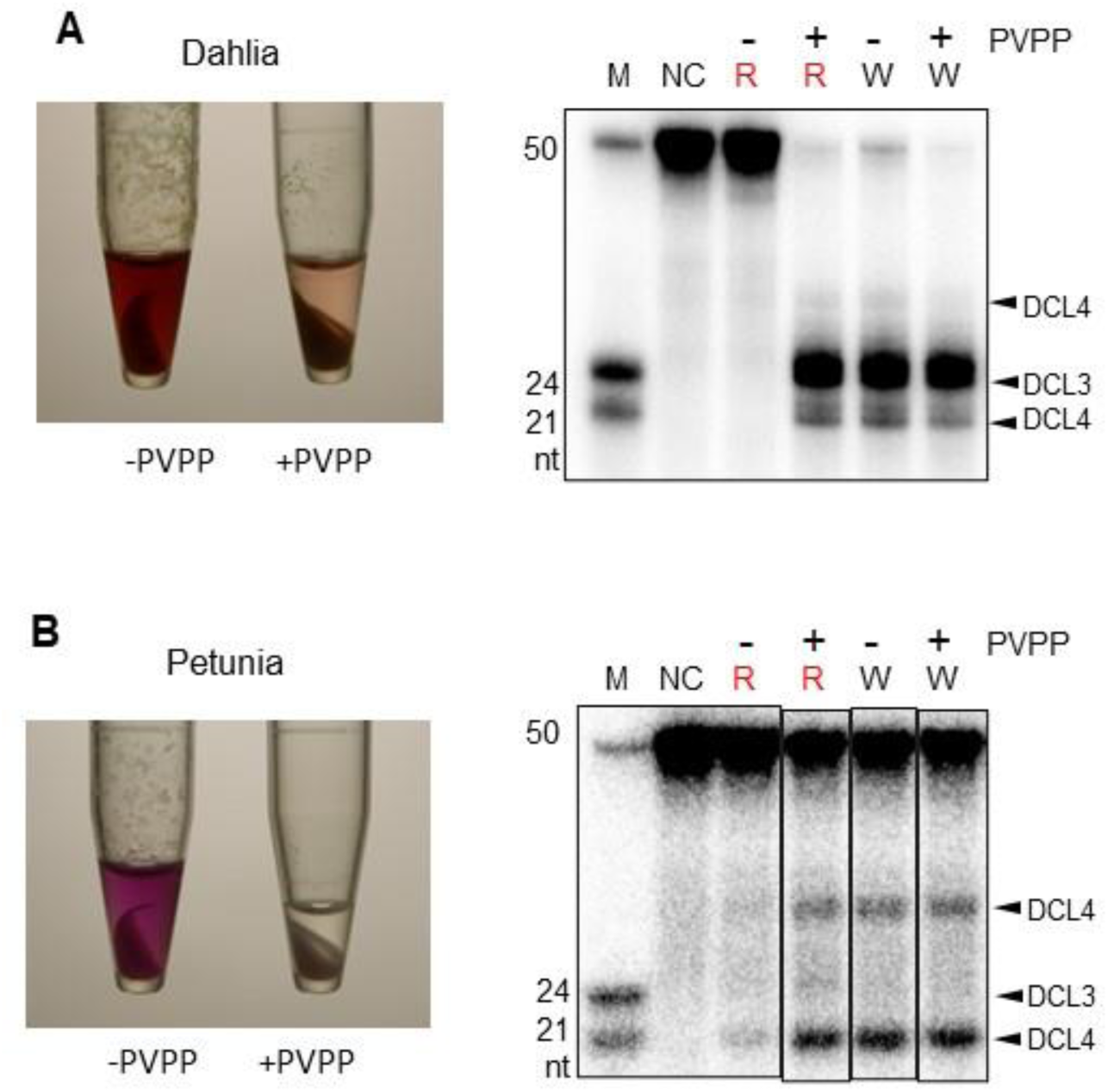
Recovery of the dicing activities of DCL3 and DCL4 by PVPP treatment. Dahlia (**A**) and petunia (**B**) petals were divided into pigmented (R) and unpigmented (W) regions, from which crude extracts were prepared. Crude extracts with (+) or without (-) PVPP were used as enzyme fractions to detect dicing activity. Photographs of centrifuged crude extracts with (+) or without (-) PVPP are shown on the left, and autoradiographs of cleaved RNA products from ^32^P-labeled 50-nt dsRNA analyzed by denaturing 15% PAGE are shown on the right. The method for detecting dicing activity is the same as that described in Figure 2. Cleaved RNA products of 21 and 31 nt were assumed to be produced by DCL4, and that of 24 nt was assumed to be produced by DCL3 (see Supplemental Figure S1). M, molecular weight markers; NC, negative control (no enzyme fraction added); R, pigmented region; W, unpigmented region: +PVPP, with PVPP treatment; -PVPP, without PVPP treatment.

### Inhibition of the dicing activities of DCL3 and DCL4 by the addition of flavonoid aglycones

To confirm that flavonoids indeed inhibit the dicing activities of DCL3 and DCL4, we selected six kinds of flavonoids (five aglycones and one glycoside) present in dahlia petals (Figure 5C), and added each of them to enzyme fractions prepared from the unpigmented region of petunia petals to perform the dicing assay. All five flavonoid aglycones inhibited dicing activities with cyanidin having the highest inhibitory activity among them (Figure 7A and Supplemental Figure S8). The concentration (155 μM) used in this experiment is a physiological concentration observed *in vivo* (Peer et al., 2001). Since both biosynthesis of flavonoid aglycones and the dicing reaction of dsRNAs into siRNAs by DCL4 occur in the cytoplasm (Hrazdina, 1992; Winkel-Shirley, 2001; Xie et al., 2004), the inhibitory interaction between flavonoid aglycones and DCL4 likely occurs *in vivo*. In contrast, cyanidin-3-glucoside showed no inhibitory activity toward DCL4, even at 1.55 mM, which is 10-fold higher than the inhibitory concentration of the cyanidin aglycone (Figure 7 and Supplemental Figures S8 and S9). Various flavonoid aglycones inhibited DCL3 and DCL4 activities, whereas the glycosylated flavonoid did not. It is well known that flavonoid aglycones are synthesized in the cytoplasm and glycosylated flavonoids, including anthocyanins, are transported from the cytoplasm into the vacuole (Hrazdina, 1992; Alfenito et al., 1998; Winkel-Shirley, 2001; Grotewold, 2006; Zhao, 2015). The addition of flavonoid aglycones to crude extracts derived from the unpigmented region of bicolor petunia petals inhibited DCL4 activity, but the treatment of these extracts with PVPP recovered DCL4 activity (Supplemental Figure S10).

**Figure 7.**
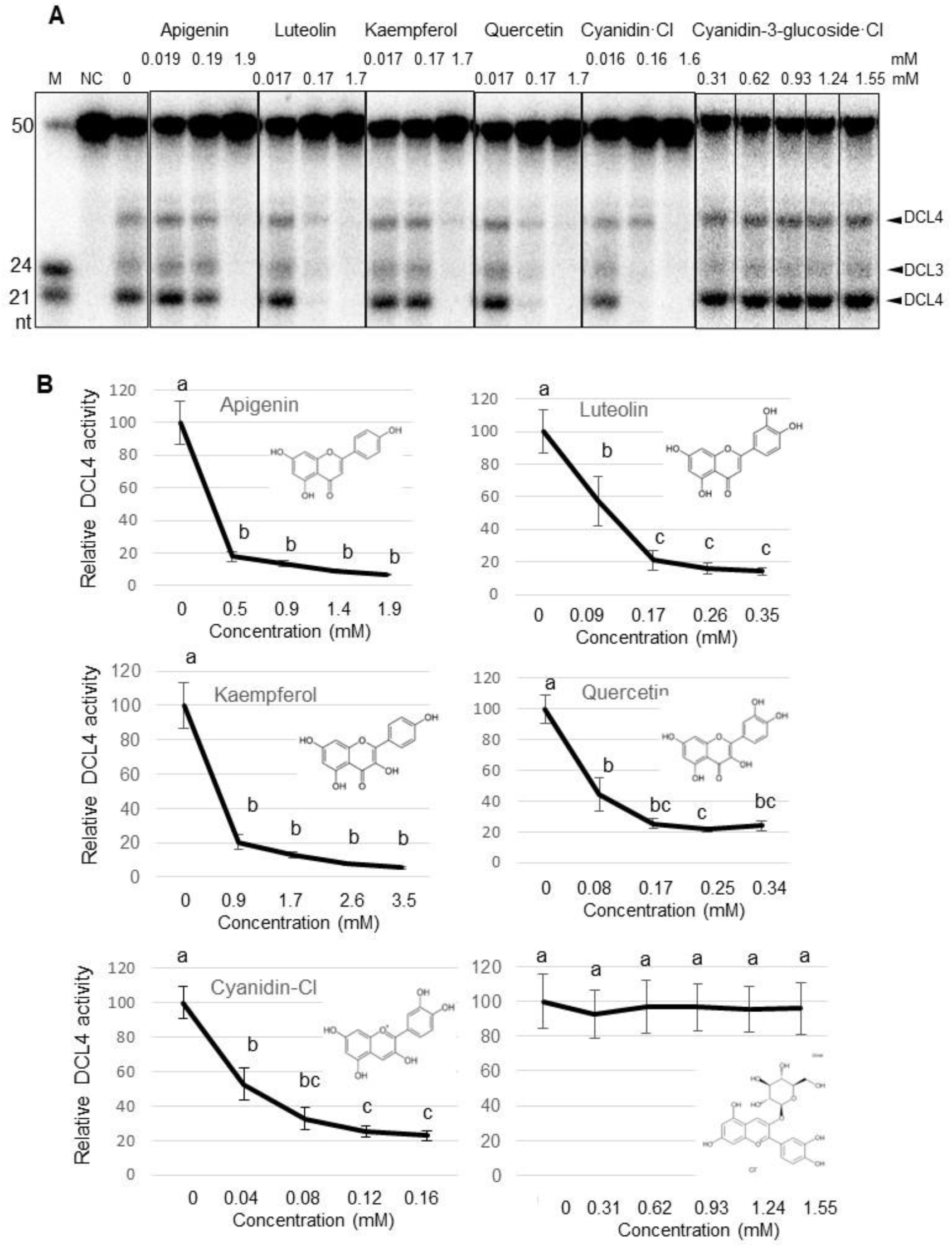
Inhibition of DCL4 activity by the addition of flavonoid aglycones. Flavonoids were added to crude extracts prepared from the unpigmented region of petunia bicolor petals and then dicing activity was analyzed. **A**) Autoradiographs of cleaved RNA products from ^32^P-labeled 50-nt dsRNA analyzed by denaturing 15% PAGE are shown. Final concentrations of flavonoids in reaction mixtures (mM) are indicated at the top of the autoradiographs. Cleaved RNA products of 21 and 31 nt were assumed to be produced by DCL4 (see Supplemental Figure S1). M, molecular weight markers; NC, negative control. **B**) Relative DCL4 activity was calculated from autoradiographs of cleaved RNA products from 50-nt dsRNA in reaction mixtures containing flavonoids as shown in (**A**). DCL4 activity in crude extracts without flavonoids was set to 100. Error bars indicate the standard errors of three biological replicates. Different letters indicate significant differences using a Tukey’s test (p<0.05).

### The dicing activity of DCL4 was detected only in the protoplasts derived from the unpigmented region of bicolor petals

The above results indicate that the flavonoid aglycones in the crude extracts prepared from the pigmented region of bicolor dahlia petals inhibited DCL4 activity (Figures 2, 4-7). However, since the enzyme fraction of the dicing assay contain both cytoplasmic and vacuolar solutes, in which secondary metabolites such as anthocyanins (glycosylated anthocyanidins) mainly accumulate, our results of the dicing activity of two DCLs might not reflect the chemical regulation of DCL activities that occurred in the cytoplasm. Therefore, we analyzed the dicing activity of intact cells by introducing ^32^P-labeled dsRNAs into protoplasts prepared from bicolor dahlia petals (Figure 8 and Supplemental Figure S11), using a protocol we established previously for *Arabidopsis* seedlings (Kakiyama et al., 2019). No cleaved RNAs were detected from colored protoplasts prepared from the pigmented region of bicolor petals, but 21-nt cleaved RNAs were detected from colorless protoplasts prepared from the unpigmented region (Figure 8 and Supplemental Figure S11). This result demonstrates that the cytoplasmic DCL4 activity producing 21-nt RNA was inhibited in the cells that constitute the pigmented region of bicolor petals. Furthermore, this result is consistent with the conclusion that the dicing activity of DCL4 is inhibited by flavonoid aglycones that cytoplasmically accumulated in the pigmented region.

**Figure 8.**
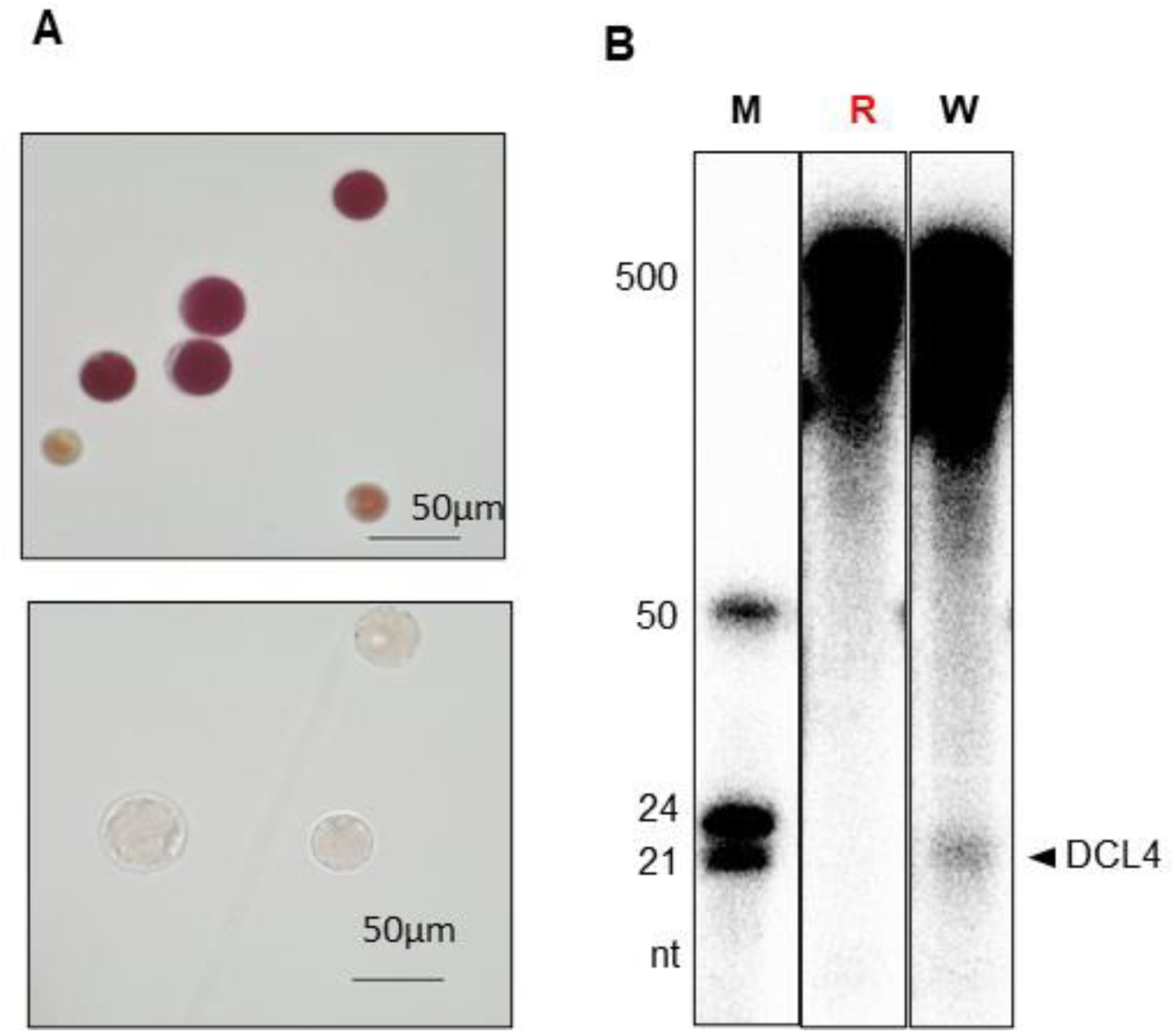
Detection of dicing activity in protoplasts prepared from the unpigmented region of bicolor dahlia petals. **A**) Photographs of protoplasts prepared from the pigmented (top) and unpigmented (bottom) regions of bicolor dahlia petals. **B**) Detection of dicing activity in protoplasts. Dahlia protoplasts were transfected with ^32^P-labeled 500-nt dsRNA by the PEG method and incubated for 1 day, and then dicing activity was detected. Cleaved product of 21-nt RNA (arrowhead) indicates DCL4 activity. R, red protoplasts prepared from the pigmented region; W, white protoplasts prepared from the unpigmented region; M, molecular weight markers.

## Discussion

Naturally occurring RNAi-mediated bicolor flowering has been reported not only in petunia (Koseki et al., 2005; Griesbach et al., 2007; Morita et al., 2012; Kasai et al., 2013) and dahlia (Ohno et al., 2011; Deguchi et al., 2013; Ohno et al., 2018) but also in gentian (Ohta et al., 2021) and bougainvillea (Ohno et al., 2021). Furthermore, in the discovery of PTGS (co-suppression) three decades ago, bicolor flowers with various patterns as well as white flowers were reported as frequently appearing in transgenic petunia plants (Napoli et al., 1990; van der Krol., 1990). However, although RNAi has been reported to be the cause of bicolor flower formation, the mechanism by which RNAi induced and uninduced regions coexist in the same petal has not been uncovered (Figure 1). In most reports of naturally occurring RNAi including in soybean cultivars with a bicolor seed coat (Cho et al., 2017), almost all target genes silenced by RNAi encode enzymes responsible for anthocyanin biosynthesis (e.g., CHS) (Napoli et al., 1990; van der Krol., 1990; Koseki et al., 2005; Ohno et al., 2011). Since CHS is a key enzyme in an initial step of the flavonoid biosynthetic pathway, suppression of CHS gene expression affects the biosynthesis of most flavonoids as well as anthocyanins (Winkel-Shirley, 2001). Therefore, we hypothesized that anthocyanins or secondary metabolites produced by the same biosynthetic pathway might be involved in the spatial regulation of RNAi. In this study, we investigated the correlation between petal pigmentation (anthocyanin accumulation) and DCL4 activity in bicolor dahlia and petunia plants (Figure 2), the accumulation of siRNAs derived from the CHS gene in the unpigmented region of petals (Figure 3), the accumulation of inhibitors and its impact on dicing activity (Figure 4), the binding of phenolic compounds to PVPP in the pigmented region (Figure 5), the recovery of DCL4 activity by removing flavonoids by PVPP (Figure 6), the ability of flavonoid aglycones to inhibit DCL4 (Figure 7), and the detection of DCL4 activity only in colorless protoplasts derived from the unpigmented region (Figure 8). In the bicolor petals of dahlia and petunia, RNAi is maintained in the unpigmented region because flavonoids that inhibit DCL4 activity are not biosynthesized there, whereas in the pigmented region, the accumulation of flavonoids that inhibit DCL4 activity inhibits RNAi, thus maintaining anthocyanin biosynthesis (Figure 9). Therefore, a clear bicolor pattern is generated by the bidirectional feedforward mechanism of antagonizing DCL4 and flavonoids (Figure 9). This study is the first to demonstrate that the bidirectional feedforward regulatory loop of DCL4 and flavonoids at the cellular level, leading to the spatial induction or uninduction of RNAi, which in turn leads to expression of the floral bicolor trait.

**Figure 9.**
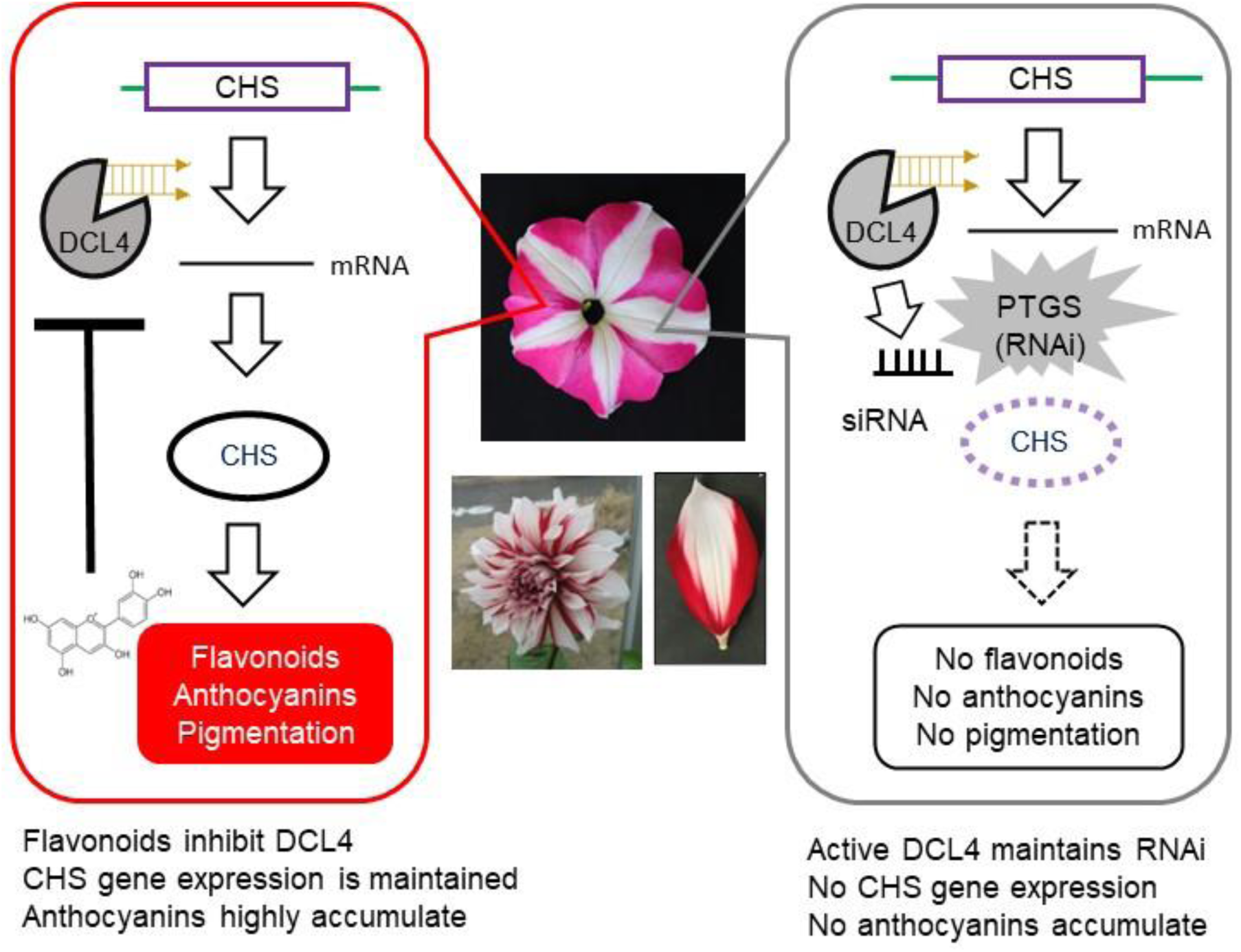
Bidirectional feedforward regulatory loop of DCL4 and flavonoids causes floral bicolor patterning in petunia and dahlia. In the pigmented (red) region, flavonoid aglycones inhibit the dicing activity of DCL4, CHS gene expression is maintained, and flavonoids including anthocyanins highly accumulate. In contrast, in the unpigmented (white) region, active DCL4 maintains RNAi (PTGS), the CHS gene cannot be expressed, and no flavonoids including anthocyanins accumulate.

Flavonoid biosynthesis takes place in the cytoplasm and flavonoids accumulate in the vacuole after glycosylation (Hrazdina, 1992; Alfenito et al., 1998; Winkel-Shirley, 2001; Grotewold, 2006; Zhao, 2015). All five flavonoid aglycones used in this study inhibited dicing activity of DCL4 (Figure 7). Conversely, cyanidin-3-glycoside did not inhibit the enzymatic activity of DCL4 (Figure 7). In other words, flavonoid aglycones localized in the cytoplasm inhibited DCL4 activity whereas the flavonoid glycoside, which accumulated in the vacuole, did not. Therefore, flavonoid aglycones may modulate DCL4 activity in the cytoplasm *in vivo*. In the dicer assay, the vacuoles are disrupted during the preparation of the crude extracts as enzyme fractions and flavonoid glycosides may be included in the enzyme fraction. However, even if flavonoid glycosides are included in the reaction mixtures, these glycosides do not inhibit DCL4 whereas flavonoid aglycones do. This conclusion is consistent with the results shown in Figure 8, in which dicing activity could not be detected in red protoplasts prepared from the pigmented regions of dahlia petals.

Flavonoids are a group of plant secondary metabolites of polyphenolic compounds that have a common chemical structure (C_6_-C_3_-C_6_), and they accumulate in various organs, such as roots, stems, leaves, flowers, and seeds, at various developmental stages (Grotewold, 2006). The major classes of flavonoids are anthocyanins (red to purple pigments), flavonols (colorless to pale yellow pigments), flavanols (colorless pigments), and proanthocyanidins or condensed tannins (Grotewold, 2006). They perform a wide range of functions, such as antioxidant activity, UV-light protection, defense against pathogens, nodulation and male fertility (Grotewold, 2006). Furthermore, flavonoids are thought to be polar auxin transport inhibitors leading to short root development (Murphy et al., 2000; Peer et al., 2004; Yin et al., 2013). Since our previous study indicated that DCL4 activity was high in meristematic tissues, such as the shoot apical meristem and floral meristem (Tabara et al., 2018), flavonoids might be involved in both auxin signaling and RNA silencing.

PVPP is used for binding and removing oxidized polyphenolic compounds, which are responsible for causing browning and bitterness in wines (Gil et al., 2017; Gil et al., 2019). It is well known that flavonoids and polyphenols function as antioxidants in cells to remove reactive oxygen species, and so they are easily oxidized. In this study, the enzymatic activity of DCL4 was recovered by PVPP to exclude flavonoids from crude extracts prepared from the pigmented regions of dahlia and petunia petals (Figure 6), and it was inhibited by the addition of flavonoid aglycones to the enzyme fraction prepared from the unpigmented region (Figure 7). The removal of flavonoids and/or oxidized flavonoids by the addition of PVPP and the addition of flavonoid aglycones might alter the redox state of the crude extracts. We have also reported that the activity of DCL4 is modulated by its redox state (Seta et al., 2017). Therefore, flavonoids might not directly interact with DCL4 to inhibit its activity, but they might affect its activity by changing the redox state of DCL4.

## Materials and methods

### Plant materials

Petunia (*P. hybrida*) and dahlia (*D. variabilis)* plants were grown on a commercially available compost soil (Akagi Engei Inc., Gunma, Japan) in a greenhouse at 24°C (Ohno et a., 2018). Commercially available seeds of petunia, cv. Rondo Rose Star were purchased from seed companies (TAKII SEED, Kyoto, Japan).

### Analysis of flavonoids and phenolic compounds

The amount of total phenolics and PVPP-bound compounds in petals was measured by using the Folin-Ciocalteu method (Makker et al., 1993; FAO/IAEA, 2000). Approximately 20 - 30 mg of petals were ground in liquid nitrogen using a mortar and pestle. Pulverized petals were suspended in 1 mL of 70% aqueous acetone, and ultrasonically treated for 20 min at room temperature by using an ultrasonic water bath (W-103T, Tokyo Garasu Kikai. Co., Tokyo, Japan). The supernatant as a total phenolic compound extract was obtained by centrifugation for 10 min at 3000g at 4°C and kept on ice. One hundred µL of this phenolic compound fraction was vigorously mixed with 100 µL of 10% (w/v) PVPP (Nacalai Tesque, Kyoto, Japan) suspension by a vortex mixer and allowed to stand for 15 min at 4°C. The supernatant as a residual phenolic compound extract to remove PVPP-bound compounds was obtained by centrifugation for 10 min at 3,000 g and 4°C. Twenty μL of the total phenolic compound extract or 50 μL of the residual phenolic compound extract to remove PVPP-bound compounds was diluted with distilled water to 305.5 μL. And 10, 20, 30, 40, and 50 μL of 0.1 mg/mL tannic acid solution, respectively, was diluted with distilled water to 305.5 μL to prepare a step dilution standard solution. Thereafter, 69.5 μL of Folin-Ciocalteu reagent (Nacalai Tesque) and 625 μL of 7.5 M sodium carbonate solution were added to these samples, respectively, and allowed to stand for 40 min after vortexing. The absorbance at 725 nm was measured. A calibration curve was obtained from the measurements of a standard sample of tannic acid, from which the amount of total phenol compounds per mg fresh weight [µg tannic acid equivalent (TAE)/mg FW] was determined using tannic acid as the standard.

The identification and quantification of flavonoids in the pigmented and unpigmented regions of bicolor petals of dahlia (cv. Yuino) and petunia (cv. Rondo Rose Star) were performed by high-performance liquid chromatography (HPLC) as described previously (Ohno et al., 2018).

### Preparation of ^32^P-labeled dsRNAs

Single-stranded RNAs (ssRNAs) 50 nt in length (Supplemental Table S1) were synthesized by Japan Bio Services Co. (Saitama, Japan), and end-labeled by T4 polynucleotide kinase (Takara Bio) and [γ-^32^P]-ATP (PerkinElmer). Equal amounts of ^32^P-labeled sense 50-nt RNA (0.01 pmol) and antisense 50-nt RNA were annealed in 10 mM Tris-HCl (pH 7.5) and 100 mM NaCl by heating at 90°C for 5 min, followed by turning off a heating block for 10 min and incubating at room temperature for 10 min (Supplemental Figure S1). A DNA template for dsRNA of 500 nt was synthesized by PCR using KOD-plus DNA polymerase (Toyobo, Osaka, Japan) with oligo DNAs containing the minimal T7-RNA polymerase promoter sequence (5′-TAATACGACTCACTATAGGG-3′) as the primers from a plasmid containing the GFP gene. ^32^P-labeled dsRNAs were synthesized using the *in vitro* T7 Transcription kit (Takara Bio) with [α-^32^P]-UTP (PerkinElmer) according to the manufacturers’ recommendations. Nucleotide sequences of ssRNAs and DNA primers used to amplify DNA templates for long dsRNA synthesis are listed in Supplemental Table S2.

### Dicer (dicing) assay using cell-free extracts

The Dicer (dicing) assay was performed as described previously (Fukudome et al., 2011; Nagano et al., 2014). Plant samples, including dahlia and petunia petals and flower buds, were collected and homogenized in 6 mL g^-1^ of extraction buffer containing 20 mM TrisHCl (pH 7.5), 4 mM MgCl_2_, 5 to 10 mM DTT, 1 mM phenyl methy sulfonyl fluoride, 1 μg mL^-1^ leupeptin, and 1 μg mL^-1^ pepstatin A at 4°C using a mortar and pestle. For the preparation of crude extracts for depleting PVPP-bound compounds, samples were homogenized with the extraction buffer containing 10% (w/v) PVPP and homogenates were placed at 4°C for 15 min. Homogenates were centrifuged twice at 20,000 *g* at 4°C for 10 to 15 min to remove debris, and the supernatant, in which protein concentration was about 12-14 mg/mL, was collected as a crude extract. In a standard dicing reaction, 1 μL of ^32^P-labeled 50-nt dsRNA with blunt ends (final concentration, ∼ 0.5 nM) was incubated with 15 μL of crude extracts and 4 μL of 5× dicing buffer containing 100 mM Tris-HCl (pH 7.5), 250 mM NaCl, 25 mM MgCl_2_, 25 mM ATP and 1.0 U μL^-1^ RNase inhibitor (Takara Bio) at 22°C for 1 to 2 h. After incubation, the cleavage products were purified by phenol/chloroform/isoamyl alcohol, separated by 15% denaturing PAGE with 8 M urea, and detected by autoradiography. Quantification of cleaved RNA products was calculated from relative band intensities measured with a Typhoon FLA 7000 image analyzer (GE Healthcare). Dicing (small RNA-producing) activity was calculated from autoradiographs as the intensity of RNA bands [21 nt (DCL4), 24 nt (DCL3) and 31 nt (DCL4)] relative to the total intensity of all bands (cleaved RNA bands and remaining substrate bands) in each lane (see Supplemental Figure S1).

### Detection of small RNAs

Approximately 5 μg of total RNA was electrophoresed in 18% denaturing polyacrylamide gels containing 1 × TBE buffer [89 mM Tris (pH 8.3), 89 mM boric acid, 2 mM EDTA] and 7 M urea, and blotted onto a nylon membrane (Zeta-Probe, Bio-Rad, Hercules, CA, USA) by electroblotting (300 mA for 1 h at room temperature). DNA fragments of the petunia *PhCHS-A* and dahlia *DvCHS2* genes as probes were amplified by PCR, and then probes for siRNA detection were made using the BcaBEST Labeling Kit (Takara Bio) with [α-^32^P]dCTP (PerkinElmer, Waltham, MA, USA). PCR primers are listed in Supplemental Table S2. Hybridization was carried out in Perfect Hyb Plus hybridization buffer (Sigma-Aldrich, St. Louis, MO, USA) containing ^32^P-labeled DNA probe at 50°C for 16 h. Membranes were washed four times in 2 × SSC (1 × SSC, 0.15 M NaCl, 15 mM sodium citrate) with 0.5% SDS at 50°C for 15 min, and then analyzed by a Typhoon FLA 7000 image analyzer (GE Healthcare, Chicago, IL., USA) (Fukuhara et al., 2011).

### Monitoring of dicing activity in protoplasts

Protoplasts were isolated from dahlia petals using a method described previously (Kakiyama et al., 2019). Approximately 2.5 g of dahlia petals was sliced with a razor blade, and 25 mL of enzyme solution [0.25 M sorbitol, 10 mM MES-KOH (pH 5.7), 10 mM CaCl_2_, 10 mM KCl, 1% cellulase Onozuka R-10, and 0.6% Macerozyme R10 and 100 μg/mL cefotaxim] was added to the sliced petals. The mixture was shaken at 30 rpm in the dark at 22°C for 4 h, the released protoplasts were sieved through a nylon mesh and Miracloth (EMD Millipore, Billerica, MA, USA) and transferred into 50-mL centrifuge tubes. Protoplasts and plant debris retained on the nylon mesh were gently sieved-through one more time by washing the mesh with 5 mL of W5 solution (0.1% glucose, 0.08% KCl, 0.9% NaCl, 1.84% CaCl_2_, and 2 mM MES-KOH [pH 5.7]). Sieved-through protoplasts were combined in a 50-mL centrifuge tube and centrifuged for 5 min at 100 g at 22°C. Protoplasts were washed twice with 15 mL of W5 solution and suspended in 1 mL of W5 solution. The concentration of protoplasts was measured by cell counting using a hemocytometer.

Protoplasts in W5 solution were incubated on ice for 30 min, after which the W5 solution was discarded, and protoplasts were resuspended in MMG solution [4 mM MES-KOH (pH5.7), 0.4 M mannitol, and 15 mM MgCl_2_] to a final concentration of 10^6^ protoplasts mL^-1^. Aliquots of protoplasts (100 µL) were transferred into a 2-mL round-bottom microcentrifuge tube and mixed gently with [α-^32^P]-UTP-labeled dsRNA of approximately 500 nt and 110 µL of PEG-calcium solution (40% PEG-4000, 0.2 M mannitol, and 100 mM CaCl_2_). Protoplasts were incubated for 7 min at room temperature. Transfection was terminated by diluting the mixture with 600 µL of W5 solution. Transfected protoplasts were collected by centrifugation for 3 min at 100 g, resuspended in 1 mL of W5 solution, and kept in the dark for 1 day. After that, total RNAs were extracted from protoplasts using phenol/chloroform/isoamyl alcohol (25:24:1), precipitated with ethanol, separated using 15% denaturing PAGE with 8 M urea, and detected by autoradiography (Typhoon FLA7000 image analyzer, GE Healthcare).

## Acknowledgements

We would like to thank Mr. Shingo Uchiya for advice on dahlia cultivation.

## Author contributions

K.K., H.M. and T.F. planned and designed the research; K.K., S.O., N.Y. and M.T. performed the experiments and analyzed the data; and K.K., S.O., H.K. and T.F. wrote the manuscript.

